# PqqU (PA2289) is responsible for Pyrroloquinoline Quinone Uptake in *Pseudomonas aeruginosa*

**DOI:** 10.64898/2026.04.27.721047

**Authors:** Christos Paschalidis, Manon Ferry, Anne-Eloïse Revillot--Schmidt, Françoise Hoegy, Gaëtan L. A. Mislin, Johana Chicher, Emmanuel Boutant, Isabelle J. Schalk, Olivier Cunrath

## Abstract

*Pseudomonas aeruginosa* relies on the redox cofactor pyrroloquinoline quinone (PQQ) for efficient glucose and ethanol metabolism via periplasmic dehydrogenases (Gcd and ExaA). While PQQ biosynthesis is well-characterized, its uptake mechanisms remain unclear. Here, we identify PA2289 (PqqU), a TonB-dependent transporter, as the primary PQQ importer in *P. aeruginosa*. Growth assays with PQQ-deficient mutants (Δ*pqqABCDEH*) demonstrated that PqqU is essential for exogenous PQQ uptake, rescuing growth on glucose and ethanol. Genomic analysis across 210 *P. aeruginosa* and 263 *Pseudomonas* strains revealed high conservation of PQQ biosynthesis and utilization genes, while PqqU showed lower prevalence (47.7%) in the genus. Transcriptional analyses using fluorescent reporters and qRT-PCR demonstrated that PqqU expression remains unchanged in response to PQQ, varying carbon sources, or iron availability, suggesting constitutive regulation. Comparative proteomics between wild-type and Δ*pqqABCDEH* strains, cultured on glucose or ethanol, uncovered extensive proteomic shifts, underscoring *P. aeruginosa’s* metabolic adaptability. Additionally, PQQ-dependent metabolic pathways appear to indirectly influence iron homeostasis, most likely through environmental acidification. Together, these results emphasize the critical role of PqqU in PQQ uptake and its broader significance in shaping the metabolic and environmental versatility of *Pseudomonas*.

## Introduction

*Pseudomonas aeruginosa*, a ubiquitous Gram-negative bacterium, is renowned for its metabolic versatility and opportunistic pathogenicity. Its ability to thrive in diverse environments is underpinned by a highly branched respiratory metabolism, enabling the utilization of a wide array of carbon sources, including glucose and ethanol (Dolan *et al*. 2020). Central to this metabolic flexibility is the redox cofactor pyrroloquinoline quinone (PQQ) (Hauge 1964) which is essential for the activity of periplasmic glucose and alcohol dehydrogenases (Gcd and ExaA, respectively) (Hommes *et al*. 1984, Görisch 2003). These enzymes catalyse the oxidation of glucose and ethanol, generating gluconate and acetaldehyde, respectively, and feeding electrons into the respiratory chain, processes critical for both energy production and environmental adaptation in *Pseudomonas spp*. (Yamada *et al*. 1993, Diehl, Wintzingerode von, and Görisch 1998). Gluconate, a product of glucose oxidation, not only serves as a carbon source but also plays a role in iron homeostasis by chelating and solubilising iron-oxide minerals (Sasnow, Wei, and Aristilde 2016), thereby facilitating its uptake and contributing to *Pseudomonas’* ability to scavenge iron in iron-limited environments.

While *P. aeruginosa* synthesizes and secretes PQQ (via its biosynthesis locus *pqqABCDEH*), the mechanisms governing its re-import remain unexplored. This is particularly relevant given that PQQ is not only a metabolic cofactor but also a potential public good in microbial communities. TonB-dependent transporters (TBDTs) are outer membrane proteins that mediate the energy-dependent uptake of scarce or large molecules, such as siderophores, vitamins, and carbohydrates, by harnessing the proton motive force via the TonB-ExbB-ExbD complex. Recent studies in *Escherichia coli* have demonstrated that PQQ can be actively imported via the TBDT PqqU, enabling the bacterium to utilize extracellular PQQ for glucose metabolism even at nanomolar concentrations (Hantke and Friz 2022, Munder *et al*. 2025).

*P. aeruginosa* PAO1 encodes a large repertoire of 35 TBDTs, yet none have been characterized for PQQ uptake to date, despite the presence of a functional PQQ biosynthesis locus (Ferry *et al*. 2024). Notably, *P. aeruginosa* encodes a putative TBDT (PA2289) located in close genomic proximity to the glucose dehydrogenase gene (*gcd*). Here we characterize the role of PA2289 in PQQ uptake and its implications for glucose and ethanol metabolism in *P. aeruginosa*, its prevalence within *P. aeruginosa* strains and more broadly within the species of *Pseudomonas*, providing insights into the interplay between metabolite secretion and re-assimilation in this versatile pathogen.

## Results

### PA2289 (PqqU) is the native transporter for PQQ in *Pseudomonas aeruginosa*

*Pseudomonas* requires the cofactor PQQ to efficiently metabolize both glucose and ethanol (Hommes *et al*. 1984, Görisch 2003). To elucidate the specific function of the TBDT PA2289 in these processes, our study first focused on glucose catabolism. We initially constructed an isogenic mutant strain lacking the ability to synthesize PQQ (Δ*pqqABCDEH*), which exhibited a marked reduction in growth fitness when glucose was provided as the sole carbon source (Fig.1). While supplementation with exogenous PQQ did not affect the growth of the wild-type strain (Fig. 1A), PQQ supplementation of approximately 10 nM fully restored growth fitness in the Δ*pqqABCDEH* mutant, confirming the specific requirement for PQQ for efficient glucose catabolism (Fig. 1D).

**Figure 1.**
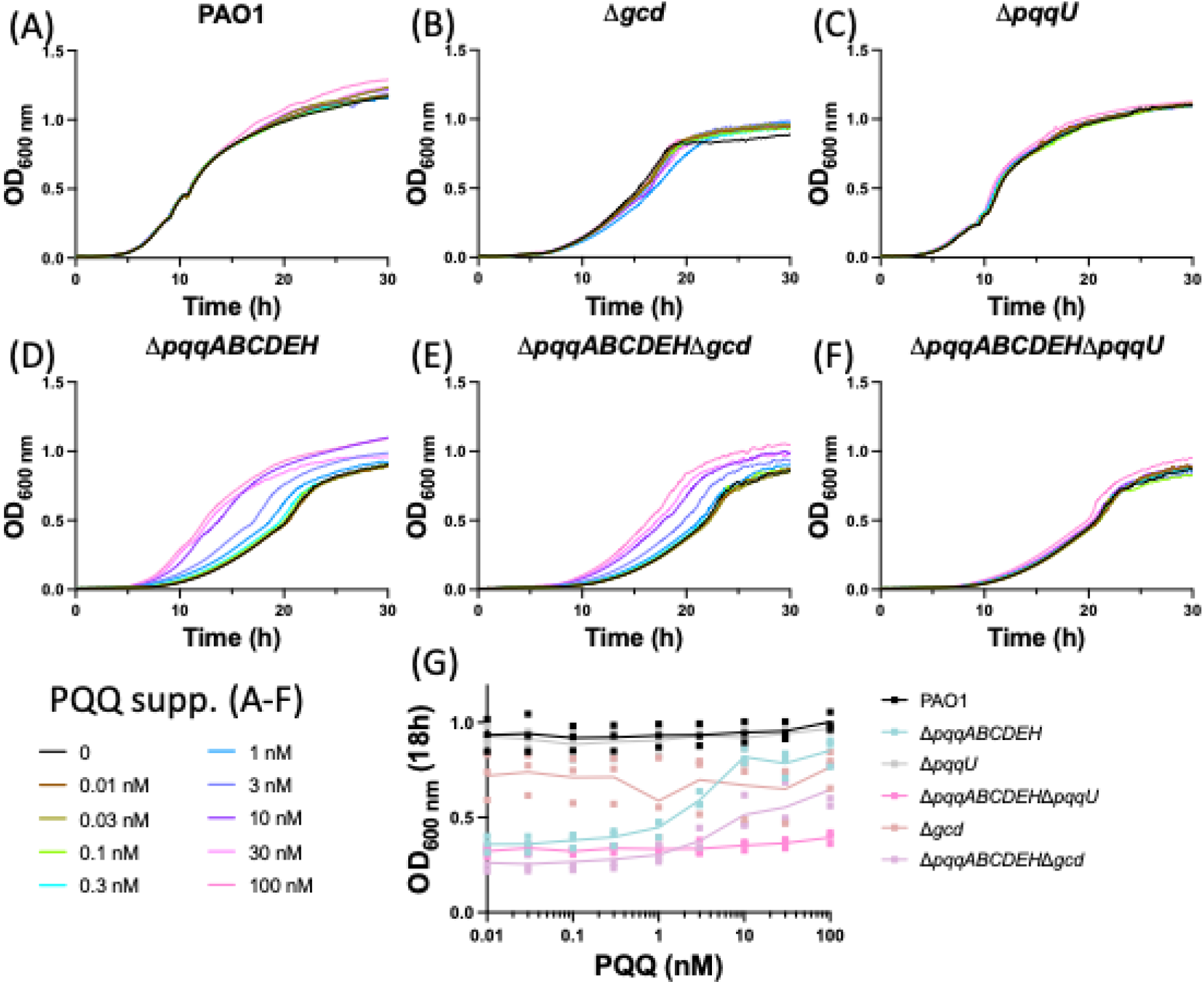
PqqU is required for exogenous PQQ uptake when grown on glucose. Bacterial strains were grown in Low-phosphate M9-Glucose with increasing concentrations of PQQ (10 pM to 100 nM) for 48 h at 37ºC and OD_600 nm_ was measured for PAO1 (**A**); Δ*gcd* (**B**); Δ*pqqU* (**C**); Δ*pqqABCDEH* (**D**); Δ*pqqABCDEH*Δ*gcd* (**E**) and Δ*pqqABCDEH*Δ*pqqU* (**F**). Mid-exponential phase OD_600 nm_ at 18 h was plotted from all conditions and all six strains (**G**).

To further explore the involvement of the periplasmic glucose dehydrogenase (Gcd), a known PQQ-dependent enzyme, we generated a Δ*gcd* mutant. This strain displayed only a slight reduction in growth fitness compared to the wild-type, but not as severe as the Δ*pqqABCDEH* mutant (Fig. 1B). The double mutant (Δ*pqqABCDEH*Δ*gcd*) exhibited a growth defect similar to that of the Δ*pqqABCDEH* single mutant, yet exogenous PQQ partially rescued its growth to levels comparable to those of the Δ*gcd* single mutant (Fig. 1E). These observations suggest the presence of an additional PQQ-dependent enzyme involved in glucose metabolism beyond Gcd.

To investigate the mechanism of PQQ uptake, we hypothesized that the TBDT PA2289 (PqqU) might facilitate PQQ transport across the outer membrane. We generated a Δ*pqqU* single mutant and a Δ*pqqABCDEH*Δ*pqqU* double mutant. The Δ*pqqU* single mutant did not show a growth defect, as it retains the ability to endogenously produce PQQ (Fig. 1C). However, the Δ*pqqABCDEH*Δ*pqqU* double mutant displayed a pronounced growth defect similar to that of the Δ*pqqABCDEH* single mutant, and, unlike the Δ*pqqABCDEH* mutant, its growth defect could not be rescued by the addition of exogenous PQQ (Fig. 1F). This result strongly indicates that PqqU is essential for PQQ transport across the outer membrane (Fig. 1G).

When *P. aeruginosa* was grown on ethanol as the sole carbon source, the supplementation of exogenous PQQ to both the wild-type strain and the single Δ*pqqU* mutant, each capable of endogenous PQQ synthesis, had no discernible impact on their growth kinetics (Fig. 2A, B). The Δ*pqqABCDEH* mutant exhibited no visible growth, which could however be fully rescued by the supplementation of exogenous PQQ, albeit at a higher concentration compared to glucose metabolism (Fig. 2C). The Δ*pqqABCDEH*Δ*pqqU* double mutant, however, failed to restore growth even at high nanomolar concentrations of PQQ, but restored growth partially only at micromolar concentrations (3 µM) (Fig. 2D, E). This phenomenon has been previously documented in *Escherichia coli* and is postulated to result from passive diffusion of PQQ through porins (Hantke and Friz 2022).

**Figure 2.**
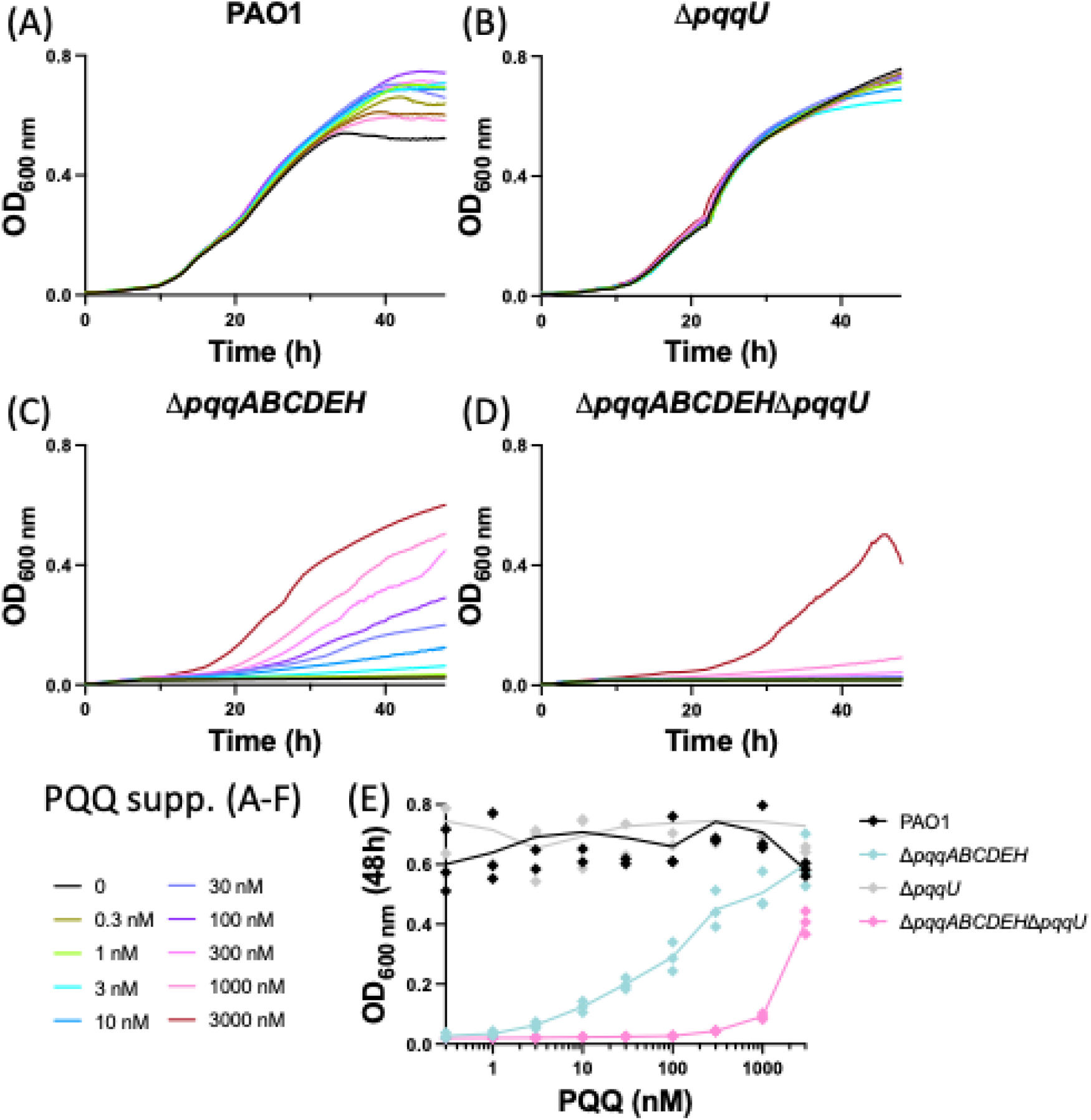
PqqU is required for exogenous PQQ uptake when grown on ethanol. Bacterial strains were grown in Low-phosphate M9-EtOH with increasing concentrations of PQQ (0.3 nM to 3000 nM) for 48 h at 37ºC and OD_600 nm_ was measured for PAO1 (**A**); Δ*pqqU* (**B**); Δ*pqqABCDEH* (**C**) and Δ*pqqABCDEH*Δ*pqqU* (**D**). Final OD_600 nm_ at 48 h was plotted from all conditions and all four strains (**E**).

### Conservation of PQQ-related genes in *P. aeruginosa* and the *Pseudomonas* genus

TBDTs such as PqqU, are widely conserved across *P. aeruginosa*, as demonstrated in one of our study (Ferry *et al*. 2024). While Munder *et al*. previously established the broad distribution of PqqU among Gram-negative bacteria (Munder *et al*. 2025), their work also revealed a marked imbalance between PQQ production and utilization, with many phyla encoding PqqU while lacking the biosynthetic machinery required for PQQ synthesis. Here, we sought to determine whether PQQ exploitation constitutes a conserved ecological strategy within the *Pseudomonas* genus and *P. aeruginosa* species. Specifically, we assessed whether the presence of the PQQ transporter PqqU is systematically associated with the PQQ biosynthetic operon (*pqqABCDEH*), or whether these functions are decoupled, consistent with the exploitation of exogenous PQQ. In parallel, we examined the conservation of PQQ-dependent dehydrogenases (ExaA and Gcd) to evaluate whether PQQ utilization is maintained independently of its endogenous production.

To address these questions, we analysed two curated datasets: 210 genomes of *P. aeruginosa* (selected to represent multilocus sequence type [MLST] diversity) and 263 genomes spanning the *Pseudomonas* genus (dereplicated at 95% average nucleotide identity [ANI]). Both datasets underwent stringent quality filtering to ensure completeness, minimal contamination, and genomic consistency (see Methods).

Our analysis identified genes with sequence homology to the PQQ biosynthetic operon (*pqqABCDEFH*), the PqqU transporter, and the PQQ-dependent dehydrogenases (*gcd* and *exaA*) across the selected genomes. Functional motifs within these homologs were verified, the conservation of these sequences within the phylogeny was analysed, and their distribution was mapped to elucidate evolutionary patterns within and beyond *P. aeruginosa*. (Fig. 3 and fig. S1).

**Figure 3.**
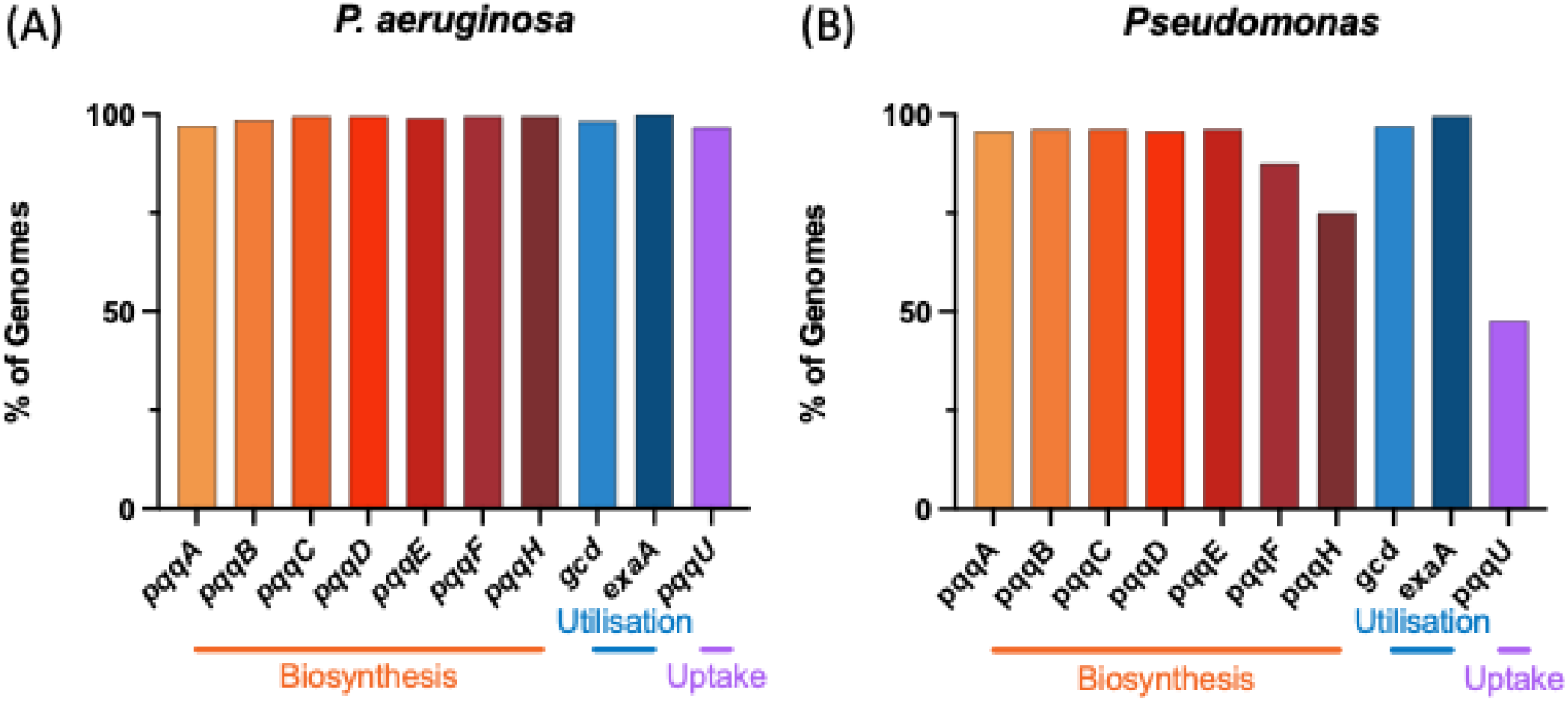
Comparative prevalence of PQQ biosynthesis, utilization and uptake genes in *P. aeruginosa* and the *Pseudomonas* genus. Bar plots representing the frequency of gene presence across two genomic datasets: (**A**) a cohort of *P. aeruginosa* (n=210) and (**B**) a representative set of the *Pseudomonas* genus (n=263). Genes include the core biosynthetic operon (*pqqABCDE*), accessory genes (*pqqF, pqqH*), the PQQ-dependent dehydrogenases (*gcd, exaA*), and the transporter *pqqU*.

Using this approach, we observed that PQQ biosynthesis (>97.1% conservation), utilization (98.1% for *gcd* and 100% for *exaA*), and uptake (96.7% for *pqqU*) were broadly conserved within *P. aeruginosa*. However, when extending our analysis to the broader *Pseudomonas* genus, we found that while the core PQQ biosynthetic genes (*pqqABCDE*, >95.8% conserved) and utilization genes (97% for *gcd* and 99.6% for *exaA*) remained highly conserved, the accessory genes (*pqqF* and *pqqH*) exhibited reduced conservation, with 87.5 and 75%, respectively. Notably, the PqqU transporter displayed unexpectedly lower conservation (47.7%) across the genus, with no discernible phylogenetic pattern (fig. S1).

### PQQ does not induce *pqqU* transcription

TBDT transcription is frequently induced by their endogenous ligands, as exemplified by siderophore transporters, which are upregulated in response to their cognate iron-chelating compounds (Ankenbauer and Quan 1994, Cobessi *et al*. 2005, Schalk and Perraud 2023). Beyond ligand-mediated regulation, certain TBDTs have their transcription and expression also modulated by environmental signals, such as nutrient availability, enabling bacteria to adapt their transport systems to fluctuating conditions (Ferry *et al*. 2024, Schalk 2025). We tested increasing concentrations of PQQ, its cognate substrate as potential regulatory ligand, as well as different culture media containing casamino acids, glucose, or ethanol as sole carbon sources (glucose and ethanol catabolism being partially or completely PQQ-dependent). Furthermore, we investigated the effect of iron availability, considering its known role in the regulation of certain TBDTs and the previously documented connection between PQQ-dependent gluconate production and iron in *P. putida* (Sasnow, Wei, and Aristilde 2016).

To assess the transcriptional response, we employed a epitopic, fluorescent transcriptional reporter (P_*pqqU*_-*ypet*) developed in a prior study (Ferry *et al*. 2024), in which the promoter region of the *pqqU* gene drives the expression of YPet, a fluorescent protein that does not interfere with the spectral properties of *P. aeruginosa’s* endogenous siderophore pyoverdine. Each construct also constitutively expresses mCherry as an internal normalization marker. Under all tested conditions, including varying PQQ concentrations, carbon sources, and iron availability, no significant changes in expression were observed (Fig. 4 and fig. S2). To further explore the potential regulatory role of PQQ, we repeated the experiments using PQQ-deficient strain Δ*pqqABCDEH*, which lacks endogenous PQQ production. However, even in this genetic background and in contrast what has been shown on the PQQ biosynthesis locus (Gliese, Khodaverdi, and Görisch 2010a), no significant *pqqU* transcription differences were detected using the fluorescent reporter system (Fig. 4).

**Figure 4.**
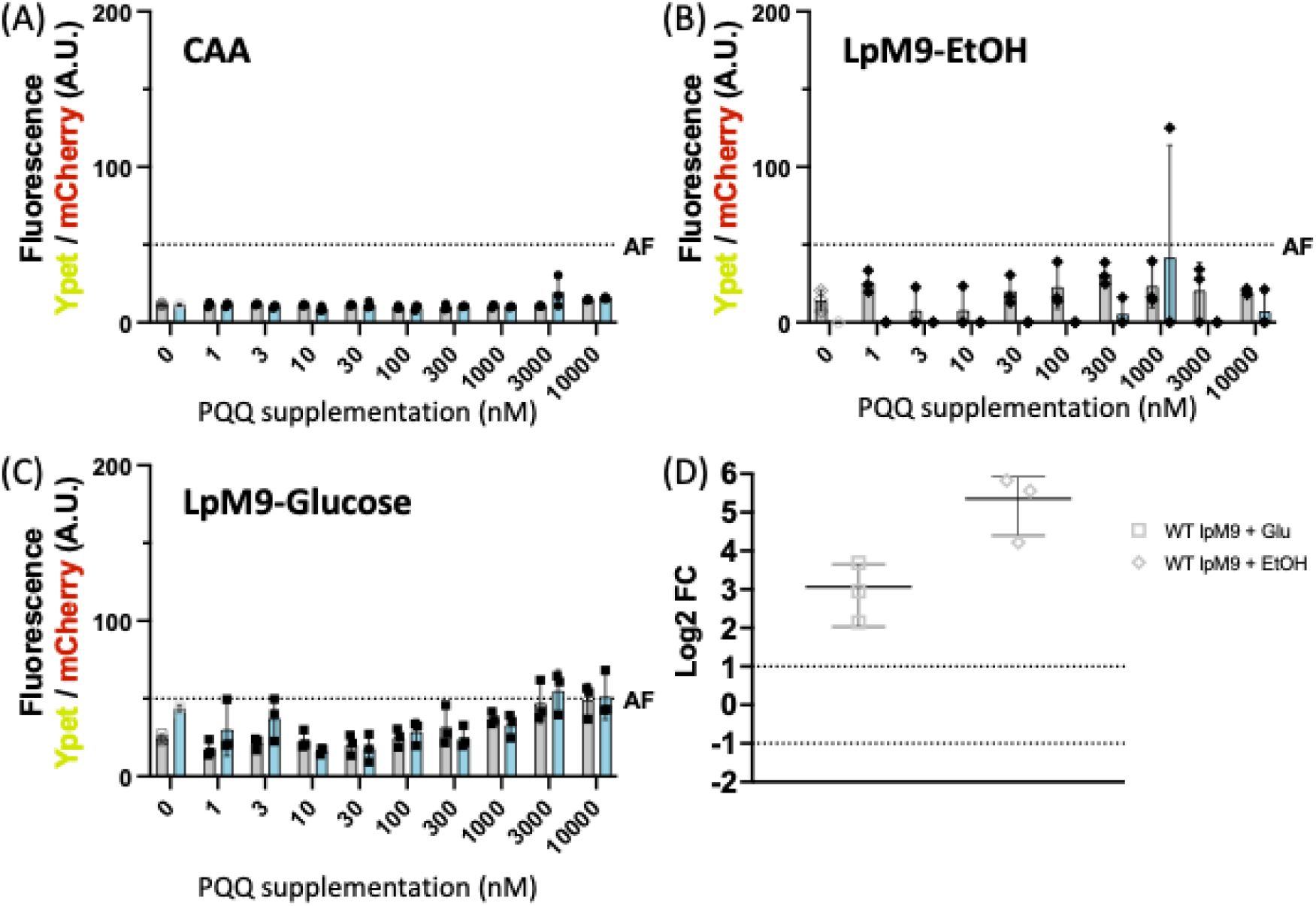
PqqU is not induced by its endogenous ligand. Bacterial strains were grown in CAA (**A**); Low-phosphate M9-EtOH (**B**) and Low-phosphate M9-Glucose (**C**) with increasing concentrations of PQQ (1 nM to 10 µM) for 24 h at 37ºC and OD_600 nm_ as well as fluorescence (mCherry and Ypet) were measured for PAO1 (grey) and the mutant Δ*pqqABCDEH* (cyan) both carrying the plasmid pOPC-273 (P_*pqqU*_-*ypet*). The inducible Ypet fluorescence was normalised to the constitutive mCherry fluorescence; AF - Autofluorescence (**D**) WT (grey) and Δ*pqqABCDEH* (cyan) cells were grown in various condition such as Low-phosphate M9 medium with glucose (square) or ethanol (diamond) as carbon source; or CAA medium (circle) with (filled) or without (clear) supplementation of 30 nM PQQ. Transcription levels of *pqqU* were quantified by qRT-PCR. The results are presented as the ratio of values obtained under the described growth conditions relative to those of the WT strain grown in CAA medium.

Given the possibility that fluorescent reporters might lack the sensitivity required to detect subtle transcriptional changes, we complemented our analysis with quantitative reverse transcription PCR (qRT-PCR), known for its higher sensitivity. Consistent with the fluorescent reporter results, even qRT-PCR analysis revealed no significant differences in the transcription levels of *pqqU*. As an internal control, we included the PQQ biosynthesis gene (*pqqC*), which exhibited significant expression variation in line with previously published data, thereby validating the robustness of our experimental approach (Magnusson *et al*. 2004, Gliese, Khodaverdi, and Görisch 2010a, Meyer *et al*. 2011). Collectively, these findings suggest that the expression of *pqqU* is not significantly influenced by PQQ availability, the carbon source, or iron levels under the conditions tested.

### PQQ dependent glucose metabolism alters *Pseudomonas’* proteome

Despite no visible transcriptional changes in *pqqU*, we wanted to further elucidate the impact of PQQ on the general physiology of *P. aeruginosa*, we conducted a comparative proteomic analysis. We examined the wild-type strain PAO1 grown in LpM9 medium with glucose as the sole carbon source, the Δ*pqqABCDEH* mutant similarly cultured in LpM9 with glucose, and PAO1 grown in LpM9 with ethanol as the carbon source.

The proteomic comparison between PAO1 and the Δ*pqqABCDEH* mutant, both grown in LpM9 with glucose, revealed substantial alterations in the proteome (fig. S3A). Notably, proteins such as ArnA, ArnB, ArnC, ArnD, MsbA, LptC, LptD, LptF, WaaA, and Wzz exhibited elevated expression in the wild-type strain, indicating differences in the outer membrane envelope and lipopolysaccharide (LPS) composition compared to the mutant. Additionally, the upregulation of PhoP, PmrA/B and PagL suggests an acidic stress response in the wild-type strain. The increased expression of alternative respiration modules, including NirS, NirF, NirM, CcoO2, CcoP1, CcpA, NosZ, and CycB, may reflect reduced oxygen availability. Elevated levels of ribosomal proteins (RpsC, RpsD, RpsG, RpsI, RpsK, RpsL, RpsM, RpsN) further support faster growth in the wild-type strain, consistent with our growth assays (Fig. 1). Conversely, the mutant displayed heightened expression of proteins involved in PQS and phenazine pathways, as evidenced by increased levels of LasI, RhlR, RsaL, PqsA, PhzA1/A2/B1/C1/M, and PhnA. These findings suggest that the absence of PQQ biosynthesis profoundly reshapes the *P. aeruginosa’s* proteome across multiple functional levels when grown in LpM9 with glucose.

To explore potential alterations in metal availability, we analysed all detected metal-binding proteins using the UniProt database Gene Ontology annotations. This approach revealed an increased abundance of magnesium-, iron-, and zinc-binding proteins in the wild-type strain when grown on glucose (fig. S3B and Table S4). Although the magnesium transporters MgtA and MgtE were undetectable, we noted elevated expression of the magnesium transporter CorA, indicating a potential increase in magnesium demand. For zinc, no changes were observed in ZnuA or ZnuC levels, nor in the pseudopaline locus (CntL/M/I or CntO); however, PA2911 protein levels (upregulated during zinc starvation) were increased. We also observed strong upregulation of the pyochelin biosynthesis pathway in the wild-type strain compared to the mutant (2–3-fold increase), while the pyoverdine pathway remained unaffected (fig. S3C). Additionally, we detected higher levels of TBDTs involved in iron uptake.

When comparing the wild-type strain grown in LpM9 with glucose versus ethanol, we observed proteomic changes similar to those described above. The glucose-grown wild-type strain exhibited increased ribosomal protein levels, an enhanced acidic stress response, elevated envelope stress markers, and reduced expression of proteins involved in PQS pathways. In contrast, growth on ethanol led to a marked upregulation of the quinoprotein ethanol system (ExaA, ExaB, ExaC) and PQQ biosynthesis (fig. S4A). Analysis of metal-binding proteins revealed only a slight increase in magnesium-binding proteins during glucose growth, with no significant changes in zinc or iron-binding protein abundance (fig. S4B) or in proteins related to pyoverdine and pyochelin production (fig. S4C).

Given the proteomic similarities between the Δ*pqqABCDEH* mutant grown on glucose and the wild-type strain grown on ethanol (fig. S5), we hypothesized that glucose-grown wild-type *P. aeruginosa* engages in a metabolic process unavailable to the other conditions, potentially the PQQ-dependent production of gluconate, as previously suggested in *P. putida*. This gluconate production may acidify the medium, explaining the observed proteomic shifts. To further investigate the link between PQQ, gluconate and iron, we investigated how gluconate, and indirectly PQQ, affects iron homeostasis in *P. aeruginosa*.

### PQQ is not directly involved in iron metabolism

To investigate a potential connection between PQQ and iron homeostasis, we first assessed siderophore production as an indicator of the iron starvation response in our experimental conditions. Specifically, we compared siderophore levels between the WT strain and the Δ*pqqABCDEH* mutant, both cultured in LpM9 medium supplemented with glucose, the main condition where we observed PQQ-dependent differences in the proteome. Proteomic analysis revealed no significant changes in the expression of proteins involved in the pyoverdine biosynthesis pathway. However, we observed a marked reduction in pyoverdine production in the WT strain grown in LpM9 with glucose, whereas the Δ*pqqABCDEH* mutant exhibited significantly higher pyoverdine levels (Fig. 5A). The same was true for pyochelin levels, which were low in the WT strain or the mutant strain supplemented with PQQ. To determine whether PQQ itself functions as a siderophore, as suggested elsewhere (Xiong *et al*. 2012, 2024), we evaluated its iron-chelating capacity using the chrome azurol sulfonate (CAS) assay. No iron complexation was detected within the testable concentration range (Fig. 5B). Higher micromolar concentrations could not be assessed due to PQQ’s intrinsic orange colour, which interfered with the colorimetric assay. To further explore PQQ’s potential role in iron acquisition, we employed a growth inhibition assay. This assay measures the ability of a given molecule to inhibit the growth in a metallophore-deficient mutant (unable to produce pyoverdine, pyochelin, or pseudopaline; Δ*pvdF*Δ*pchA*Δ*cntL*) that additionally lack all three *tonB* genes (Δ*tonB1*Δ*tonB2*Δ*tonB3*) which makes it unable to take up siderophore-complexed iron. If the tested molecule chelates iron and would act as an iron-chelator, this mutant is unable to grow as all iron is locked away for its growth. While the addition of the known iron-chelating siderophore desferoxamine completely inhibits the mutants’ growth, supplementation with 10 µM PQQ did not, reinforcing the conclusion that PQQ does not act as a siderophore per se (Fig. 5C). To elucidate the relationship between PQQ and iron homeostasis, we next examined gluconate production, a PQQ-dependent process, and its impact on environmental pH. In the presence of PQQ, *P. aeruginosa* WT – PAO1 strain acidified the medium significantly, an effect absent in the Δ*pqqABCDEH* mutant strain unless exogenous PQQ was added (Fig. 5D). Notably, medium acidification, rather than gluconate levels (Fig. 5E), correlated with reduced pyoverdine production (Fig. 5F). When the WT strain was cultured in buffered LpM9 or when the Δ*pqqABCDEH* mutant was supplemented with PQQ in buffered conditions, gluconate production remained high, though pyoverdine levels were sustained. These findings suggest that pyoverdine biosynthesis may be downregulated due to acidification rather than gluconate levels in the environment, linking PQQ levels indirectly to iron homeostasis.

**Figure 5.**
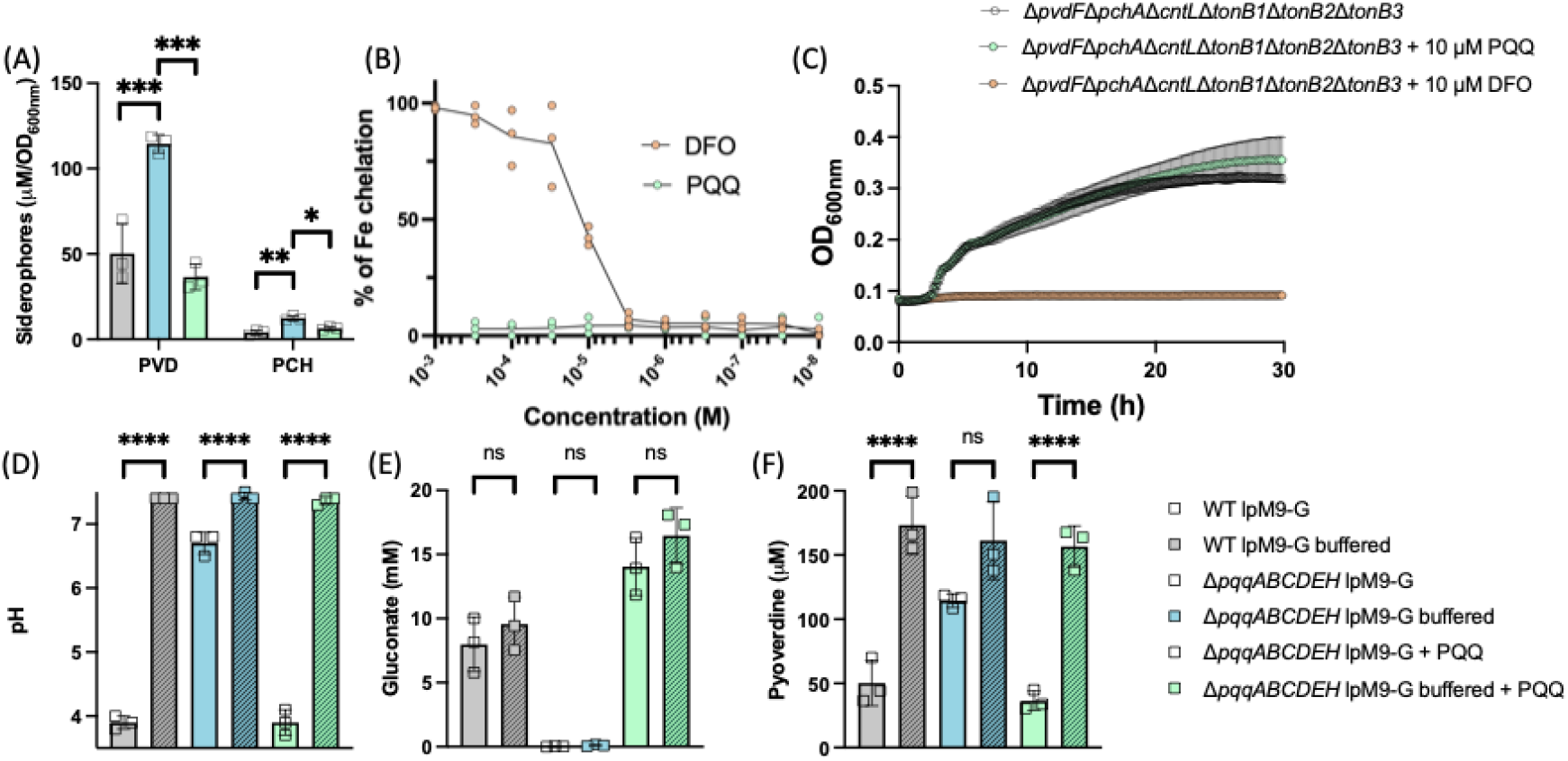
PQQ is not directly linked to iron homeostasis. (**A**) Cells were grown in Low phosphate M9 glucose medium for 16 h and siderophore concentration was measured subsequently. (**B**) CAS assay using both PQQ and a known siderophore DFO. (**C**) Growth kinetics of *P. aeruginosa* mutant, unable to produce its endogenous siderophores in the iron restricted medium CAA, alone (clear circles) with the addition of 1 μM of DFO-B (orange) or 1 μM of PQQ. (**D-F**) Cells were grown in Low phosphate M9 glucose unbuffered (clear squares) or buffered (filled squares) medium for 16 h and pH (**D**), gluconate concentration (**E**) and pyoverdine concentration (**F**) were measured.

## Discussion

Our study identifies PqqU – PA2289 as the first experimentally validated non-metal-dependent TBDT in *P. aeruginosa*. Unlike most TBDTs, which are associated with the uptake of metals, or metal-bound ligands such as siderophores (Ferry et al., 2024; Schalk, 2025), PqqU functions independently of metal availability. This finding highlights the need for further investigation into the ligands, regulatory mechanisms, and environmental cues governing non-metal-dependent TBDTs, such as Starch Utilization Systems (Sus) found in *Bacteroides* or *Porphyromonas* (Glenwright *et al*. 2017, Madej *et al*. 2020, Gray *et al*. 2021, Pollet *et al*. 2023), which may play underappreciated roles in *P. aeruginosa’s* bacterial metabolism, adaptation, and virulence. We demonstrate that PqqU is the primary transporter for PQQ uptake in *P. aeruginosa*. The inability of the Δ*pqqABCDEH*Δ*pqqU* double mutant to rescue growth on glucose or ethanol with exogenous PQQ, unlike the Δ*pqqABCDEH* single mutantconfirms PqqU’s essential role in PQQ import at nanomolar concentrations. However, partial growth rescue at micromolar PQQ concentrations suggests that one of the 41 porins (Ude *et al*. 2021) may facilitate passive PQQ uptake under these conditions, as previously suggested in *Escherichia coli* (Hantke and Friz, 2022; Munder et al., 2025).

Our bioinformatic screen suggests that PqqU exhibits lower conservation across the *Pseudomonas* genus (47.7%), with no clear phylogenetic pattern, while being highly conserved within *P. aeruginosa* (96.7%) (Fig. 3). This raises questions about alternative PQQ uptake mechanisms or niche-specific adaptations in some *Pseudomonas* species. The presence of PqqU in some Gram-negative bacteria (Munder *et al*. 2025) does not imply universal conservation across all members of a given genus. Indeed, our findings reveal variable distribution of PqqU within the *Pseudomonas* genus, where many species lack this transporter. This variability highlights the importance of investigating the conservation patterns and ecological roles of TBDTs across diverse bacterial lineages. The secretion and re-import of PQQ may reflect cooperative dynamics in microbial communities, akin to other public goods like siderophores or vitamin B12 (Guadarrama-Orozco *et al*. 2023). While only a limited number of bacterial species synthesize PQQ (Munder *et al*. 2025), a broad range of bacteria, including non-producers, can uptake and utilize PQQ via transporters like PqqU and PQQ-dependent enzymes such as Gcd. This widespread capacity to exploit PQQ may drive its importance in producing strains, reinforcing the need for efficient and rapid uptake mechanisms. PqqU appears to be particularly important at nanomolar PQQ concentrations (Fig. 1 and 2), suggesting an exceptionally high affinity for its ligand. To date, only the affinities of ferri-siderophores for their respective TBDTs have been extensively studied in *P. aeruginosa*, typically falling in the low nanomolar range (Schalk, Mislin, and Brillet 2012, Moynié *et al*. 2019, Schalk 2025). Given PqqU’s functional significance at nanomolar PQQ levels, it is plausible that its affinity for PQQ is similarly high, if not even stronger. Finally, PqqU expression appears low and constitutive, as it is not induced by PQQ, carbon sources, or iron availability, as confirmed by fluorescent reporters and qRT-PCR. This contrasts with the regulation of many –but not all– siderophore transporters and suggests alternative regulatory mechanisms for PqqU (Schalk 2025).

The reduced conservation in the *Pseudomonas* genus of the accessory genes *pqqF*, 87.5% (encoded on a separate gene locus in PAO1) and *pqqH*, 75% suggests potential functional redundancy or niche-specific adaptations. In contrast, the high conservation (>97%) of the PQQ core biosynthesis enzymes (*pqqABCDE*) and utilization genes (*gcd* and *exaA*) in *P. aeruginosa* underscores their essential role in central metabolism. Additionally, the partial growth rescue of the Δ*pqqABCDEH*Δ*gcd* mutant with exogenous PQQ (Fig. 1F) implies the existence of additional, uncharacterized PQQ-dependent pathways beyond Gcd, warranting further investigation. In *P. aeruginosa*, only Gcd and ExaA have so far been identified as PQQ□dependent, whereas PQQ□dependent dehydrogenases for mannitol, glycerol, sorbitol and methanol have been described in other bacteria (Goodwin and Anthony 1998, Adachi *et al*. 2003, Hölscher and Görisch 2006, Gliese, Khodaverdi, and Görisch 2010b). Additional studies are required to determine whether homologs of these enzymes or others exist in *P. aeruginosa* and could account for the partial rescue observed here.

Proteomic analyses revealed profound shifts in the Δ*pqqABCDEH* mutant, including upregulation of outer membrane and LPS biosynthesis proteins, as well as stress response pathways (e.g., PhoP, PmrA/B). This highlights PQQ’s role in shaping *Pseudomonas’* metabolism, which in turn can affect envelope integrity and adaptive responses. PQQ-dependent glucose metabolism in *Pseudomonas* leads to the accumulation of gluconate and subsequent acidification of the extracellular environment, conferring several ecological advantages to the producing organism. This acidification not only inhibits the growth of pH-sensitive competitors (Kaur *et al*. 2006) but also enhances the solubilization of essential nutrients such as phosphate and iron, a phenomenon previously documented in *P. putida* (Sasnow, Wei, and Aristilde 2016). By converting glucose, a widely metabolizable carbon source, into gluconate, which is less accessible to many competing microorganisms, *Pseudomonas* employs a competitive strategy termed “metabolic niche modification” or “resource amending” (Odum and Barrett 1971, Hibbing *et al*. 2010, Sasnow, Wei, and Aristilde 2016). This strategy is not unique to *Pseudomonas* but represents a broader microbial behaviour observed in diverse bacterial species. Prominent examples include the conversion of lactate to acetate by lactic acid bacteria (Vásquez-Dean *et al*. 2026) and the sequestration of iron through siderophore secretion (Niehus *et al*. 2017, Figueiredo *et al*. 2022), as seen in *P. aeruginosa* and other species. Such metabolic adaptations provide *Pseudomonas* with a distinct competitive advantage during interbacterial competition, enabling it to thrive in complex and nutrient-limited environments. This ability to reshape its ecological niche underscores the organism’s versatility and dominance in microbial communities.

While Xiong *et al*. suggested that in *Salmonella* YncD (now called PqqU) may be involved in iron homeostasis, via siderophore uptake (Xiong *et al*. 2012, 2024), Hantke and Friz already concluded that PQQ may not act as a siderophore (Hantke and Friz 2022). While we show again that PQQ does not function as a siderophore, here confirmed by CAS assays and growth rescue experiments, which demonstrate its lack of iron-chelating activity at physiological concentrations, its metabolic byproduct, gluconate, indirectly modulates iron homeostasis. The acidification mediated by gluconate production, increases iron’s solubility and drastically reduces the biosynthesis of the endogenous siderophores pyoverdine and pyochelin, a response driven by pH-dependent regulatory mechanisms rather than direct iron chelation. This phenomenon aligns with observations in *E. coli* and *Salmonella*, where acidification similarly reduces siderophore production, suggesting a conserved bacterial trait (Valdebenito *et al*. 2006, Ferry *et al*. 2026). Although gluconate possesses iron-chelating potential at high concentrations, its involvement in iron acquisition, akin to iron-citrate uptake, warrants further investigation.

## Conclusion

The metabolic versatility conferred by PQQ-dependent pathways enables *P. aeruginosa* to thrive across diverse environments, contributing to its versatility. While targeting PqqU or PQQ biosynthesis could disrupt metabolic adaptability, its therapeutic potential may be limited, as PqqU loss results in only minor fitness defects. Given that all here tested *P. aeruginosa* strains, including clinical isolates, possess PqqU, it may be regarded as potential gateway for antimicrobial delivery, using the Trojan-horse strategy. However, its low expression levels might pose significant challenges for efficient drug delivery. Finally, future work should explore the broader ecological and mechanistic implications of PqqU in *P. aeruginosa* pathogenicity and focus on the identification of non-metal dependent TBDTs more broadly.

## Acknowledgments

We are indebted to the members of the MMBCA team for discussion.

## Author contributions

Conceptualisation: OC. Methodology: MF, CP and OC. Investigation: MF, FH, CP, AERS, FH, GLAM, JC, EB, OC. Visualisation: MF, CP and OC. Funding acquisition: IJS, OC. Supervision: IJS, OC. Writing – original draft: OC. Writing – review and editing: MF, CP, IJS and OC.

## Data and materials availability

Mass spectrometry data are available via ProteomeXchange with identifier PXD077532. All data are available in the main text or supplementary materials.

## Materials and Methods

### Chemicals

Pyrroloquinoline quinone (PQQ), was obtained from MedChemExpress (HY-100196) and was solubilised in MilliQ water at 3 mM and was kept for maximum 6 months at – 20 ºC.

### Bacterial Strains and Growth Conditions

*P. aeruginosa* strains used are listed and described in **Table S1** in the Supporting Information. Bacteria were routinely grown in Lysogeny broth (**LB**) (AthenaES Ref:0102) at 37 ºC, shaking. PQQ growth experiments were performed in Low phosphate M9 (**LpM9**) medium at 37 ºC, shaking – 1.28 g/L Na_2_HPO_4_ (Merck Ref: 231-448-7), 0.3 g/L KH_2_PO_4_ (VWR Ref: 26936.260), 0.5 g/L NaCl (Euromedex Ref: 1112-C), 1 g/L NH_4_Cl (Merck Ref: 1145), 10 µM of FeCl_3_ (Prolab, Ref: 24 207.291) and 10 mL/L Goodiemix. <u>Goodiemix solution</u> consists of 94.89 g/L MgSO_4_ 7H_2_O (Merck Ref: 1.05886.1000), 1.11 g/L CaCl_2_ (Sigma-Aldrich Ref: 31307), 0.033727 g/L thiamine hydrochloride (Sigma-Aldrich Ref: T4625-10G) and 125 mL/L trace element solution [13.08 g/L MgCl_2_ 6H_2_O (Merck Ref: 1.05833.1000), 2 g/L CaCl_2_ (Sigma-Aldrich Ref: 31307), 0.90 g/L ZnSO_4_ (Strem chemicals Ref: 93-3045), 0.85 g/L MnSO_4_ · H_2_O (Bio Basic Ref: MB0334), 0.24 g/L CuSO_4_ · 5 H_2_O (Strem Ref: 50-901-14907), 0.06 g/L H_3_BO_3_ (Sigma-Aldrich Ref: B0394-500G), 51 mL/L 1M HCl (Thermo-Fischer Ref: H/1200/PB15)]. Either Glucose (Euromedex Ref: UG3050) or Ethanol (Carlos Erba, Ref: 524125) were used as carbon source for a final concentration of 0.4 % (4 g/L or 4 mL/L, respectively). For iron-limitation we used the Casamino Acid Medium (**CAA**) at 37 ºC, shaking – 5 g/L Bacto™ Casamino acid (BD Ref: 223050), 1.46 g/L K_2_HPO_4_ 3H_2_O (Carlo Erba Ref: 471767), 0.25 g/L MgSO_4_ 7H_2_O (Merck Ref: 1.05886.1000).

### Plasmids and Genetic engineering

Gene deletions were generated as described in Hmelo et al. (Hmelo *et al*. 2015). Briefly, 700 base pairs upstream and downstream of region to be deleted were PCR amplified (Phusion^®^, NEB) and inserted into the suicide vector pEXG2, which was linearized using the restriction enzymes BamHI and EcoRI, using the NEBuilder^®^ HiFi DNA Assembly Master Mix (NEB #E2621L). The plasmid was introduced into an *E. coli* strain (TOP10). After 6h of mating between the plasmid-containing donor *E. coli* strain, the *E. coli* helper strain and the recipient *P. aeruginosa* strain, trans-conjugants were selected on LB plates containing 30 µg/mL gentamicin and 10 µg/mL chloramphenicol. Counter-selection was performed on no-salt LB plates supplemented with 10% of sucrose at 30ºC. Mutants were screened by colony PCR using described primers and sequence was verified using Sanger sequencing (Eurofins). Primers used for genetic engineering in this study are listed in **Table S2**. All plasmids used in this study are listed in **Table S3**.

### Quantitative Real-Time PCR

The transcription of selected genes was assessed using quantitative reverse transcription PCR (qRT-PCR), as previously described (Gasser *et al*. 2016, Perraud *et al*. 2020). Mid-exponentially growing bacteria (OD_600 nm_ = 0.6), 2.5 × 10L cells were harvested, and two volumes of RNAprotect Bacteria Reagent (Qiagen) were added. Total RNA was extracted from the pellet using the RNeasy Plus Mini Kit (Qiagen) following the manufacturer’s instructions. Subsequently, 1 μg of total RNA was reverse transcribed using the RNA-to-cDNA Kit (Applied Biosystems). Specific cDNA was quantified using a StepOne Plus instrument (Applied Biosystems) with the appropriate primers (**Table S2**) and Power SYBR Green PCR Master Mix (Applied Biosystems). Transcript levels for each gene were normalized to *clpx* and *rpoD* and expressed as logL fold changes relative to the reference condition.

### Bacterial growth kinetics

For glucose and ethanol growth experiments, *P. aeruginosa* PAO1 WT or mutant strains were precultured overnight in LpM9, 0.4 % glucose at 37 ºC, shaking. For iron-limited conditions, bacterial cells were precultured overnight in CAA medium supplemented with 1 µM of FeCl_3_. Overnight cultures were harvested and OD_600 nm_ was measured using an Eppendorf BioPhotometer™. OD_600 nm_ was adjusted to final OD_600 nm_ of 0.01. 96 well plates (Greiner Bio-One #655161) were prepared containing the described medium with various concentrations of PQQ. 96 well plates were incubated in a Tecan plate reader (Series Infinite M200PRO), measuring OD_600 nm_ every 15 minutes for 24 h. Data were plotted using GraphPad Prism 10.

### Fluorescence reporter measurements

*P. aeruginosa* carrying a low copy number plasmid with various P_*pqqU*_-*ypet* were precultured in CAA supplemented with 10 µM FeCl_3_ and 30 µg/mL gentamicin overnight at 37ºC, shaking at 200 rpm. Overnight cultures were diluted 1/100 to inoculate freshly prepared microplates, which contained the growth conditions described in main text supplemented with 30 µg/mL gentamicin. Plates were incubated for 20 h at 37ºC, shaking at 600 rpm. After 20 h incubation, the microplates were centrifuged using a microplate centrifuge (Sigma Laborzentrifugen™ 10155) at 3500 rpm for 10 min at room temperature. Supernatant were removed and bacterial pellets were resuspended in PBS. At least three independent replicates have been performed. Optical density (600 nm), Ypet fluorescence (ex: 500 nm, em: 540 nm) and mCherry fluorescence (ex: 570 nm, em: 610 nm) were measured using a TECAN plate reader (Series Infinite M200PRO). Data were plotted and analyzed using One-way ANOVA using GraphPad Prism 10.

### Proteomics analysis

For mass spectrometry analysis, 10µg of Pseudomonas aeruginosa proteins were precipitated twice with 0.1M ammonium acetate in 100% methanol (5 volumes, - 20°C) to get rid of detergents and other contaminants. They were reduced (5 mM Dithiothreitol, 95°C, 10min), alkylated (10mM Iodoacetamide, room temperature, 20 min) and digested overnight with 300 ng of sequencing-grade trypsin (Promega). Generated peptides were analysed by nanoLC-MS/MS on a reversed phase nanoElute 2 coupled to a TIMS-TOF Pro 2 mass spectrometer (Bruker Daltonik Gmbh) using a data-independent acquisition (DIA) strategy. Peptides were separated on the integrated emitter column IonOpticks Aurora Elite (25 cm × 75 μM, 1.7 μm particle size and 120 Å pore size; AUR3– 15075C18-CSI) with a 65 min gradient.

Data were searched using DIANN 2.0 software with a library free approach against the Uniprot database with PAO1 Pseudomonas aeruginosa taxonomy (v.2025_02, 5646 sequences) and a database containing common contaminants. Prostar software (1.38.1) was used for the statistical analyses of the intensities. Imputation of missing values was based on the 1% det quantile for both Partially Observed Values (POV) and Missing in Entire Condition values (MEC). A LIMMA statistical test and a Benjamini-Hochberg correction were used to generate log2(FC) and adjusted p-values. The mass spectrometry proteomics data have been deposited to the ProteomeXchange Consortium via the PRIDE (Perez-Riverol *et al*. 2025) partner repository with the dataset identifier PXD077532.

### Chrome azurol S

CAS solution was prepared as described by Schwyn and Neilands (Schwyn and Neilands 1987). All reagents (Chrome Azurol S, HDTMA, anhydrous piperazine and 5-sulfosalicylic acid) were purchased from Sigma-Aldrich. Chemicals were mixed with 100 µL of CAS solution in a well of a 96-well microplate with flat bottoms (Greiner Bio-One #655161). Controls included MilliQ water mixed with CAS solution and an iron-free variant of the CAS solution (APO-CAS). The plate was then incubated in a dark environment at room temperature for 1 hour and 30 minutes. The optical density at 630 nm (OD_630 nm_) was measured using a TECAN plate reader (Series Infinite M200PRO). To determine the iron chelating capability of the bacterial supernatants the following formula was used:

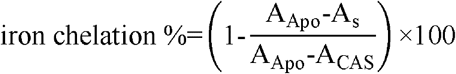

AApo: absorbance of uninoculated media + APO-CAS (positive control)

ACAS: absorbance of uninoculated media + CAS (negative control)

As: absorbance of sample

### Siderophore quantification

*P. aeruginosa* cultures were grown in various media (50 mL volumes), and supernatants were collected following centrifugation at 10,000 × g for 10 minutes. Pyoverdine quantification was performed by measuring absorbance at 400 nm when pH was around 7, using an extinction coefficient of ε 19,000 L mol□^1^ cm□^1^. In conditions where pH was low (around 4) we measured absorbance at 380 nm, using an extinction coefficient of ε 16,500 L mol□^1^ cm□^1^ (Albrecht-Gary *et al*. 1994). For pyochelin quantification, supernatants were first acidified to pH 2–3 with 2 mL of 10% citric acid, followed by one extraction with 50 mL of dichloromethane. The organic phase was dried using 25 g anhydrous Na_2_SO_4_, cotton filtered, and evaporated to dryness. Residues were then resuspended in 2 mL of methanol, and absorbance spectra were recorded from 230–500 nm in a quartz cuvette. Pyochelin concentrations were determined using the characteristic absorbance peak at 313 nm and an extinction coefficient of ε 4,900 L mol□^1^ cm□^1^.

### Bioinformatic analysis

#### Genome Selection and Quality Control

Two parallel analyses were conducted to assess gene conservation: one within *Pseudomonas aeruginosa* species and another across the *Pseudomonas* genus. Genomes were retrieved from the BV-BRC database (Wattam *et al*. 2024). Initial quality assessment was performed using CheckM statistics provided by BV-BRC. Genomes with CheckM completeness ≤ 90% or contamination ≥ 5% were excluded from further analysis.To ensure genomic consistency, the number of coding sequences (CDS) was plotted against genome length for each strain. Genomes deviating from the expected linear relationship by >500 CDS were removed (Horesh et al., 2021; Sharp and Foster, 2024). Only genomes with a minimum size of 4 million base pairs were retained for downstream analysis.

#### Genome Dataset Curation

For the *P. aeruginosa*-specific analysis, diversity was captured by selecting one representative strain per unique multilocus sequence type (MLST), while retaining the reference strains PAO1 (Genome ID: 208964.12), PA7 (Genome ID: 652611.49), and PA14 (Genome ID: 381754.6). This resulted in a dataset of 210 genomes. For the genus-level analysis, genomes were dereplicated using skani (Shaw and Yu 2023) to select one representative per cluster at 95% average nucleotide identity (ANI), yielding a dataset of 263 genomes.

#### Homolog Identification and Phylogenomic Analysis

The curated genome datasets were screened for homologs of PQQ biosynthetic genes (*pqqA, pqqB, pqqC, pqqD, pqqE, pqqF, pqqH*), the PQQ transporter *pqqU*, and PQQ-dependent dehydrogenases (*gcd* and *exaA*). The homolog identification pipeline was adapted from (Munder *et al*. 2025). The PQQ biosynthetic operon genes (*pqqABCDEFH*) were identified using cblaster (v1.3.18), with *P. aeruginosa* PAO1 sequences as queries. For the detection of PqqU homologs among *Pseudomonas* species, BLASTp (e-value ≤ 1×10□□) was performed using the PA2289 (PAO1) sequence as the query. Resulting hits were aligned using FAMSA (v1.2.5) (Deorowicz, Debudaj-Grabysz, and Gudyś 2016). Sequences were manually verified for the conserved molecular signature R304 and R365 (Munder *et al*. 2025) using Geneious (Kearse *et al*. 2012). Gcd and ExaA homologs were identified using a two-step HMMER (v3.3) (Eddy 1992) pipeline that combines target-specific profile HMMs with verification against PQQ Pfam domains (Mistry *et al*. 2021) (PF01011, PF13360, and PF13570) with an e-value ≤ 1×10□□, following established structural criteria (Diamond *et al*. 2019, Munder *et al*. 2025). For the within-species analysis (n=210 P. aeruginosa genomes), reference HMMs were built from single PAO1 sequences (Gcd: PA2290; ExaA: PA1982). For the genus-level analysis (n=263 Pseudomonas genomes), reference HMMs were built from multiple sequence alignments of three orthologs each, spanning the major Pseudomonas clades: *P. aeruginosa* PAO1, *P. putida* (BV-BRC ID 160488.4), and *P. fluorescens* (BV-BRC ID 216595.4). Sequences were aligned with FAMSA (v1.2.5), and HMMs were built using hmmbuild (HMMER v3.4) with default parameters. Each predicted proteome was searched against the specific HMMs and against Pfam profile PF01011 (the canonical PQQ enzyme repeat) using hmmsearch with an e-value cutoff of 1×10□□. A protein was assigned as Gcd or ExaA only if it returned a significant hit against both the corresponding specific HMM and PF01011. Where a single protein scored against both Gcd and ExaA HMMs, the assignment with the lower e-value was retained. Phylogenomic trees were constructed using GToTree and visualized and annotated using iTOL (v7.0) (Letunic and Bork 2021).

## Supplementary figures

**Fig. S1.**
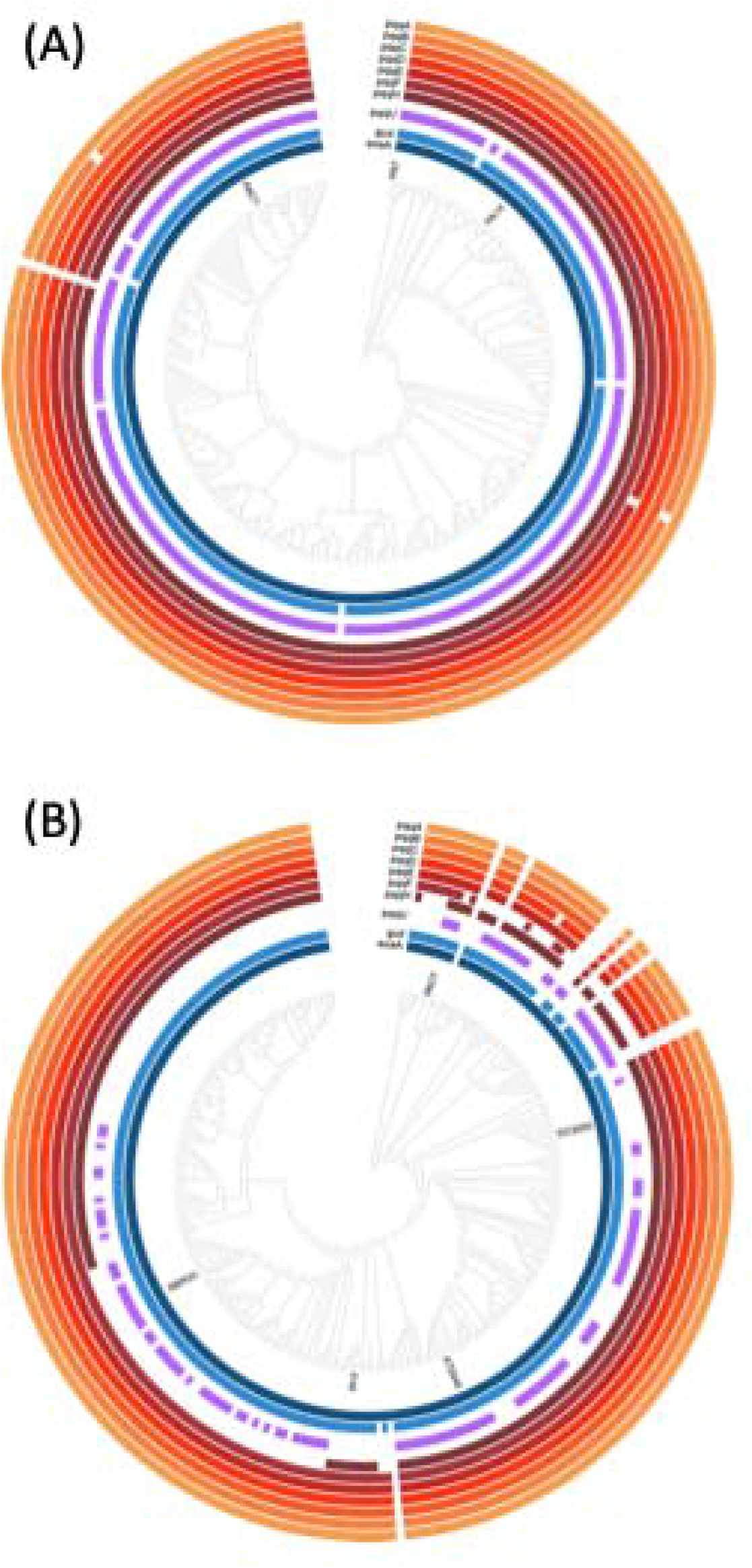
Conservation of PQQ-related genes across *P. aeruginosa* species (A) and *Pseudomonas* genus (B) highlights their ecological importance. Cladogram of 210 *P. aeruginosa* and 263 *Pseudomonas* strains, based on the concatenated alignment of universal core genes shared across all strains. Strains were selected to maximize genetic diversity with each strain having a unique MLST profile. Coloured rings depict the presence or absence of different PQQ-related genes.

**Fig. S2.**
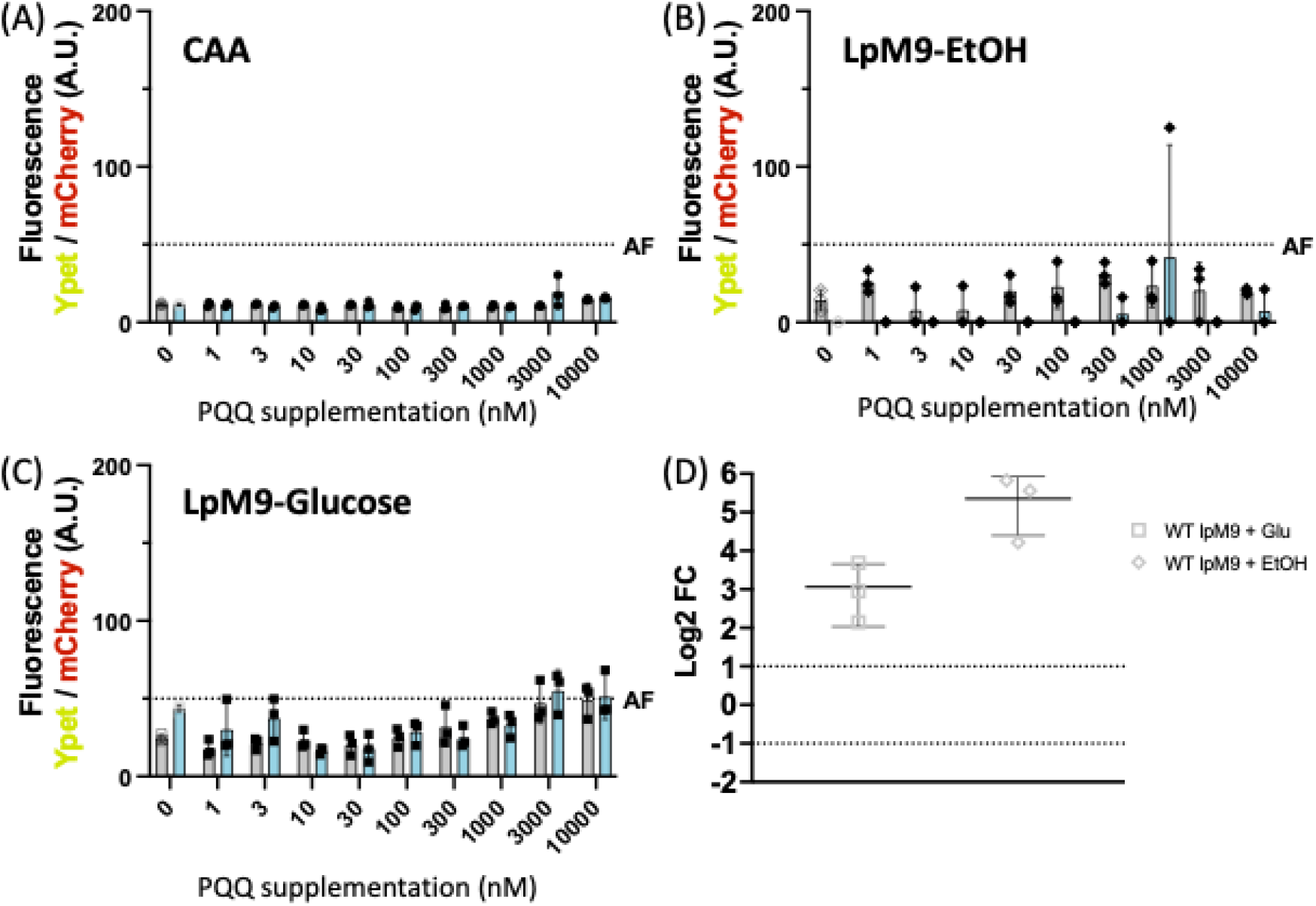
PQQ does not induce PqqU. Cells were grown in CAA + 10 µM of FeCl_3_ (**A**); LpM9-EtOH + 10 µM of FeCl_3_ (**B**) and LpM9-Glucose + 10 µM of FeCl_3_ (**C**) with increasing concentrations of PQQ (1 nM to 10 µM) for 24 h at 37ºC and OD_600 nm_ as well as fluorescence (mCherry and Ypet) were measured for PAO1 (grey) and Δ*pqqABCDEH* (cyan) both carrying the inducible plasmid pOPC-273 (P_*pqqU*_-*ypet*-P_*pilM*_-*mcherry*). The inducible Ypet fluorescence was normalised to the constitutive mCherry fluorescence; (**D**) WT – PAO1 (grey) cells were grown in various condition such as Low phosphate M9 medium with glucose (square) or ethanol (diamond) as carbon source. Transcription levels of *pqqC* were quantified by qRT-PCR. The results are presented as the ratio of values obtained under the described growth conditions relative to those of the WT – PAO1 strain grown in CAA medium.

**Fig. S3.**
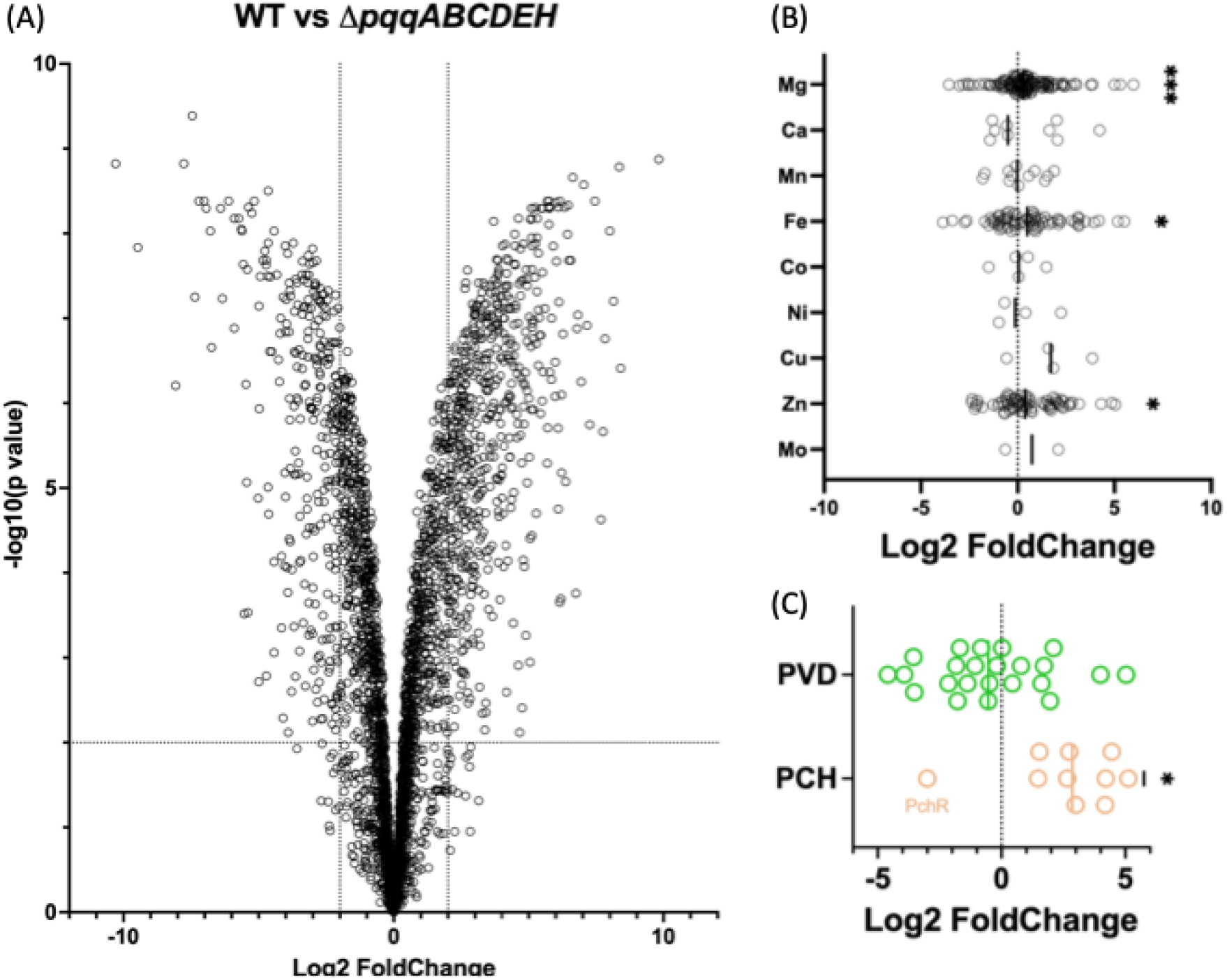
PQQ induce profound proteomic shifts. (**A**) Proteome comparison of WT and Δ*pqqABCDEH P. aeruginosa* cells grown in Low phosphate M9 medium with glucose as sole carbon source. (**B**) Metalloprotein comparison between above mentioned conditions according to their metal cofactor. (**C**) Comparison of proteins involved in pyoverdine (green) and pyochelin (brown) biosynthesis.

**Fig. S4.**
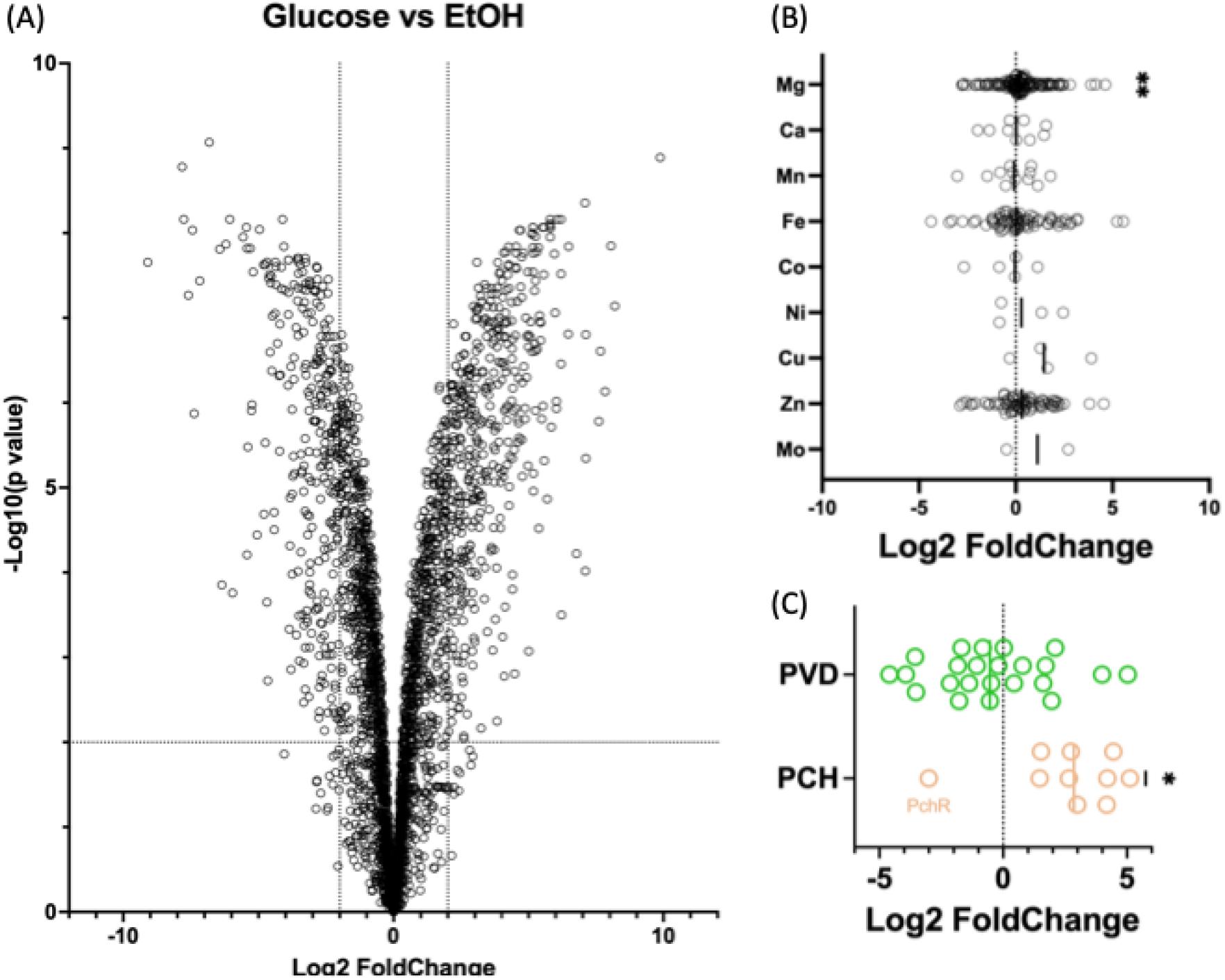
Glucose versus ethanol metabolism induce similar profound proteomic shifts. (**A**) Proteome comparison of WT *P. aeruginosa* cells grown in Low phosphate M9 medium either with glucose or ethanol as sole carbon source. (**B**) Metalloprotein comparison between above mentioned conditions according to their metal cofactor. (**C**) Comparison of proteins involved in pyoverdine (green) and pyochelin (brown) biosynthesis.

**Fig. S5.**
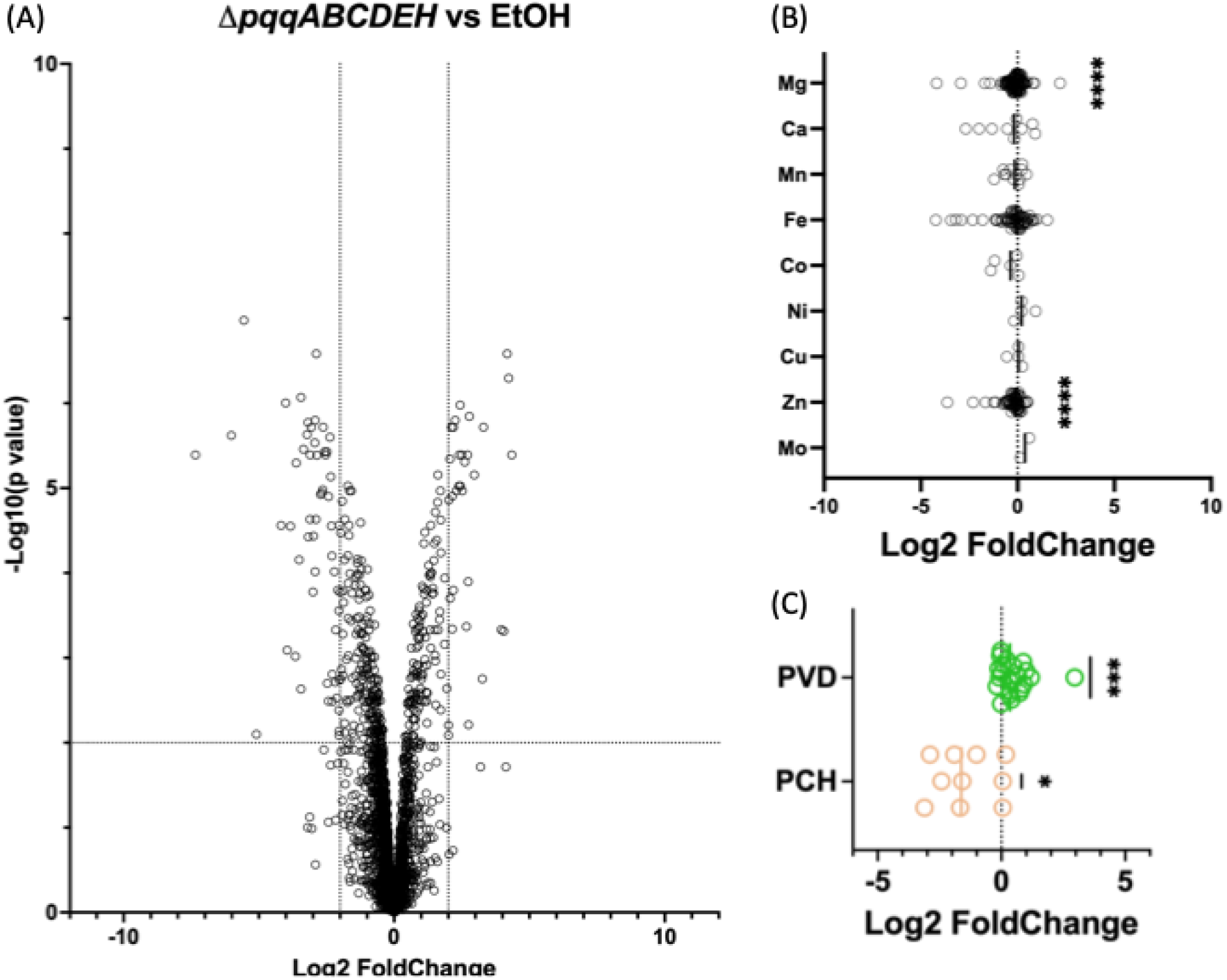
Ethanol metabolism and none-PQQ glucose metabolism have similar proteomic signatures. (**A**) Proteome comparison of WT *P. aeruginosa* cells grown in Low phosphate M9 medium either with ethanol as sole carbon source and Δ*pqqABCDEH P. aeruginosa* cells grown in Low phosphate M9 medium with glucose as sole carbon source. (**B**) Metalloprotein comparison between above mentioned conditions according to their metal cofactor. (**C**) Comparison of proteins involved in pyoverdine (green) and pyochelin (brown) biosynthesis.

