## Supplemental Table 1, 2 and 3 for "PqqU (PA2289) is responsible for Pyrroloquinoline Quinone Uptake in *Pseudomonas aeruginosa*"

**Table S1**

Bacterial strains used in this study.

| ***Escherichia coli*** | | |
| --- | --- | --- |
| **Strain** | **Description** | **Source** |
| TOP10 | F- *mcrA* Δ(*mrr*-*hsdRMS*-*mcrBC*) φ80lacZΔ*M15* Δ*lacX74* *nupG* *recA1* *araD*139 Δ(*ara-leu*)7697 *galE*15 *galK*16 *rpsL*(StrR) *endA1 λ* | Invitrogen |
| HB101 + pRK2013 | F–, *thi-1, hsdS20 (rB–, mB–), supE44, recA13, ara-14, leuB6, proA2, lacY1, galK2, rpsL20 (strr), xyl-5, mtl-1* + pRK2013 | Promega, (Figurski and Helinski 1979) |
| ***Pseudomonas aeruginosa*** | | |
| **Strain** | **Description** | **Source** |
| PAO1 | DSMZ 22644 – wild type strain | (Stover *et al.* 2000) |
| Δ*pqqABCDEH* | PAO1– lacking PQQ-biosynthesis | This study |
| Δ*gcd* | PAO1– lacking glucose dehydrogenases Gcd | This study |
| Δ*pqqABCDEH*Δ*gcd* | PAO1– lacking PQQ-biosynthesis and glucose dehydrogenases Gcd | This study |
| Δ*pqqU* | PAO1– lacking PA2289 – PQQ uptake transporter | This study |
| Δ*pqqABCDEH*Δ*pqqU* | PAO1– lacking PQQ-biosynthesis and PA2289 – PQQ uptake transporter | This study |
| Δ*pvdF*Δ*pchA*Δ*cntL*Δ*tonB1*Δ*tonB2*Δ*tonB3* | PAO1 – pyoverdine negative, pyochelin negative, pseudopaline negative – lacking all genes coding for tonB – Δ*tonB1*, Δ*tonB2* and Δ*tonB3* | This study |

**Table S2**

Primers used in this study.

| **Primer name** | **Sequence** | **Description** |
| --- | --- | --- |
| oOPC-1253 | gcataaatgtaaagcaagcttctgcaggtcgactctagagtgttccaggaggaaatcttcgg | amplify 700-*pqqABCDEH* for pqqABCDEH deletion in PAO1 |
| oOPC-1254 | ccctttcaatcctccagcaggggcttggtccacatgg |  |
| oOPC-1255 | accccatgtggaccaagcccctgctggaggattgaaaggg | amplify 700+*pqqABCDEH* for pqqABCDEH deletion in PAO1 |
| oOPC-1256 | gcacgatcatgcgcacccgtggaaattaattaaggtaccgcagtgcatcgatgccggtacaggccg |  |
| oOPC-1257 | tcaaggggcacaacggcatg | check *pqqABCDEH* deletion in PAO1 |
| oOPC-1258 | aacgcctcgatggcgtgtac |  |
| oOPC-1247 | gcataaatgtaaagcaagcttctgcaggtcgactctagagtcgctgccgacggtggct | amplify 700-*gcd* for pqqABCDEH deletion in PAO1 |
| oOPC-1248 | ggacgatgagcgaaggtaacctgcccaggcaatgacc |  |
| oOPC-1249 | gcgggtcattgcctgggcaggttaccttcgctcatcgtccggta | amplify 700+*gcd* for pqqABCDEH deletion in PAO1 |
| oOPC-1250 | gcacgatcatgcgcacccgtggaaattaattaaggtaccgggcgaaggccgatgacg |  |
| oOPC-1251 | gtacaggtggttgcggctcag | check *gcd* deletion in PAO1 |
| oOPC-1252 | gcgagtaccggctgggttac |  |
| oOPC-1160 | gcataaatgtaaagcaagcttctgcaggtcgactctagaggatccctggtgatcgtcagcggctacacc | amplify 700-*tonB1* for pqqABCDEH deletion in PAO1 |
| oOPC-1161 | ccctcatgtcgccacagcctgagaagcgccgctgaggcggttcgc |  |
| oOPC-1144 | ccgcctcagcggcgcttctcaggctgtggcgacatgag | amplify 700+*tonB1* for pqqABCDEH deletion in PAO1 |
| oOPC-1145 | gcacgatcatgcgcacccgtggaaattaattaaggtaccgaattcctggtagtggtgctggatg |  |
| oOPC-1162 | tggtggggagcgttcttcctg | check *tonB1* deletion in PAO1 |
| oOPC-1147 | aaacagctgcaggctcctcc |  |
| oOPC-1148 | gcataaatgtaaagcaagcttctgcaggtcgactctagagcatgccggggcacatgaacg | amplify 700-*tonB2* for pqqABCDEH deletion in PAO1 |
| oOPC-1149 | cagattcagttcaacttgaactggggtgttgccatgaaaag |  |
| oOPC-1150 | ttttcatggcaacaccccagttcaagttgaactgaatctgcagcgt | amplify 700+*tonB2* for pqqABCDEH deletion in PAO1 |
| oOPC-1151 | gcacgatcatgcgcacccgtggaaattaattaaggtaccggatgacgcggaacgcg |  |
| oOPC-1152 | gaagttcgccatccacccgg | check *tonB2* deletion in PAO1 |
| oOPC-1153 | ctccttcacgttgcgcgagg |  |
| oOPC-1154 | gcataaatgtaaagcaagcttctgcaggtcgactctagaggcgacgccgatctgcc | amplify 700-*tonB3* for pqqABCDEH deletion in PAO1 |
| oOPC-1155 | gcggagtgcgtcatagacttctgtccagcaagtagcgcc |  |
| oOPC-1156 | cggcgctacttgctggacagaagtctatgacgcactccgc | amplify 700+*tonB3* for pqqABCDEH deletion in PAO1 |
| oOPC-1157 | gcacgatcatgcgcacccgtggaaattaattaaggtaccgcgcagagcctgcgcg |  |
| oOPC-1158 | gcgccttcaatacgcacagc | check *tonB3* deletion in PAO1 |
| oOPC-1159 | ctacctgctggaacaggccg |  |
| *pqqU* F | CTACCAGAACTTCAACGGCG | check by qPCR *pqqU* transcription in PAO1 |
| *pqqU* R | CAGGTAGGGATCGAGGCTG |  |
| *pqqC* F | CTGGAGCACTATCGGACCC | check by qPCR *pqqC* transcription in PAO1 |
| *pqqC* R | ACGTCCAGCTTGAATTGCAG |  |
| *clpX* F | CTGCGCTCATGCAGATCCT | check by qPCR *clpX* transcription in PAO1 (control) |
| *clpX* R | TCGAACAGCTTGGCGTACTG |  |
| *rpoD* F | ACAAGATCCGCAAGGTACTGAAG | check by qPCR *rpoD* transcription in PAO1 (control) |
| *rpoD* R | CGCCCAGGTGCGAATC |  |

**Table S3**

Plasmids used in this study. Resistance: Gentamicin (*P. aeruginosa* - 30μg/mL; *E. coli* - 15μg/mL)

| **Plasmid name** | **Relevant genotype** | **Resistance** | **Source** |
| --- | --- | --- | --- |
| pEXG2 | *ori* ColE1, *rop*, *bom*, *oriT*, *sacB*, *GmR* | Gentamicin | (Rietsch *et al.* 2005) |
| pOPC-297 | pEXG2-derived suicide vector to delete *pqqABCDEH* in *P. aeruginosa* | Gentamicin | This study |
| pOPC-296 | pEXG2-derived suicide vector to delete *gcd* in *P. aeruginosa* | Gentamicin | This study |
| pVN14 | pEXG2-derived suicide vector to delete *pqqU* in *P. aeruginosa* | Gentamicin | This study |
| pOPC-281 | pEXG2-derived suicide vector to delete *tonB1* in *P. aeruginosa* | Gentamicin | (Volck *et al*. 2026) |
| pOPC-282 | pEXG2-derived suicide vector to delete *tonB2* in *P. aeruginosa* | Gentamicin | This study |
| pOPC-283 | pEXG2-derived suicide vector to delete *tonB3* in *P. aeruginosa* | Gentamicin | This study |
| pOPC-273 | pOPC-240 derived reporter plasmid with P*_pqqU_*-ypet (*PA2289*) | Gentamicin | (Ferry *et al.* 2024) |

Ferry M, Ferriz H, Sharp C *et al.* Identification of iron and zinc responsive TonB-dependent transporters in Pseudomonas aeruginosa’s. Preprint, bioRxiv, Oct. 2024, 2024.10.14.618150. https://doi.org/10.1101/2024.10.14.618150.

Figurski DH, Helinski DR. Replication of an origin-containing derivative of plasmid RK2 dependent on a plasmid function provided in trans. *Proc Natl Acad Sci U S A* 1979;**76**(4):1648–52. https://doi.org/10.1073/pnas.76.4.1648.

Rietsch A, Vallet-Gely I, Dove SL *et al.* ExsE, a secreted regulator of type III secretion genes in Pseudomonas aeruginosa. *Proc Natl Acad Sci U S A* 2005;**102**(22):8006–11. https://doi.org/10.1073/pnas.0503005102.

Stover CK, Pham XQ, Erwin AL *et al.* Complete genome sequence of Pseudomonas aeruginosa PAO1, an opportunistic pathogen. *Nature* 2000;**406**(6799):959–64. https://doi.org/10.1038/35023079.
